# Computational investigation of protein surface polarity and pH-dependence: application to antibodies shows that antibody-antigen interfaces tend to be relatively polar

**DOI:** 10.1101/345231

**Authors:** Max Hebditch, Jim Warwicker

## Abstract

Protein instability leads to reversible self-association and irreversible aggregation which is a major concern for developing new biopharmaceutical leads. Protein solution behaviour is dictated by the physicochemical properties of the protein and the solution. Optimising protein solutions through experimental screens and targeted protein engineering can be a difficult and time consuming process. Here, we describe development of the protein-sol web server, which was previously restricted to protein solubility prediction from amino acid sequence. Tools are presented for calculating and mapping patches of hydrophobicity and charge on the protein surface. In addition, predictions of folded state stability and net charge are displayed as a heatmap for a range of pH and ionic strength conditions. Tools are evaluated in the context of antibodies, their fragments and interactions. Surprisingly, antibody-antigen interfaces are, on average, at least as polar as Fab surfaces. This benchmarking process provides the user with thresholds with which to assess non-polar surface patches, and possible solubility implications, in proteins of interest. Stability heatmaps compare favourably with experimental data for CH2 and CH3 domains. Display and quantification of surface polarity and pH / ionic strength dependence will be useful generally for investigation of protein biophysics.

## Introduction

Protein biopharmaceuticals (biologics), and in particular monoclonal antibodies, are crucial for many new generation therapeutic interventions (Carter, 2011; Ecker *et al*., 2015). Compared to traditional small chemical drugs, antibodies have a higher specificity, as well as target selectivity, leading to fewer off-target effects (Smith, 2014). However, due to the liquid formulation requirements, and the general instability of proteins compared to small molecules (Wang, 1999; Manning *et al*., 2010), the development of monoclonal antibody biopharmaceuticals can be difficult. Instability of monoclonal antibody products is exacerbated by the delivery requirements. Most biopharmaceutical antibodies are delivered subcutaneously (Narasimhan *et al*., 2012), and this limits the maximum volume to around < 1.5 ml, which generally necessitates a concentration of around 100g/L or higher. This requirement further complicates the delivery of a stable protein formulation, as high concentration often leads to a less stable protein product. Protein instability can lead to non-specific association causing aberrant solution behaviours (Woods and Nesta, 2010; Liu *et al*., 2005; Raut and Kalonia, 2016), in more severe cases, instability gives rise to the formation of irreversible, and immunogenic aggregates (Hansel *et al*., 2010). Reversible and irreversible association processes limit protein solubility, and have therefore complicated the manufacturing of protein biopharmaceuticals (Shire, 2009; Daugherty and Mrsny, 2006; Mitragotri *et al*., 2014). To improve the stability and developability of biologics, various groups have focused on predicting the physicochemical properties of proteins in an attempt to accelerate drug production. Previous work within our group has looked at protein features related to protein solubility, in particular the lack of positively charged surface patches (Chan *et al*., 2013), the ratio of lysine to arginine residues (Warwicker *et al*., 2014), and the stability of individual Fab domains (Hebditch *et al*., 2017b). Experimental studies (Chari *et al*., 2009; Esfandiary *et al*., 2015; Neergaard *et al*., 2013; Yearley *et al*., 2013; Calero-Rubio *et al*., 2017; Roberts *et al*., 2014a; Ghosh *et al*., 2016; Inouye *et al*., 2016; Schermeyer *et al*., 2017), as well as computational approaches (Calero-Rubio *et al*., 2016; Lilyestrom *et al*., 2013; Corbett *et al*., 2017; Kuhn *et al*., 2017), have been aimed at understanding the solution behaviour of proteins and biologics.

Much research has focused on the role of anisotropic surface patches of charge and hydrophobicity in causing reversible and irreversible protein association (Yadav *et al*., 2012; Perchiacca *et al*., 2012; Chow *et al*., 2016; Li *et al*., 2016; Roberts *et al*., 2014b; Austerberry *et al*., 2017). This experimental work has led to efforts in predicting protein surface patches *in silico*. For example, the commercially available spatial aggregation propensity (SAP) software (Chennamsetty *et al*., 2009b) which has been applied to IgG antibodies (Chennamsetty *et al*., 2009a), and is used for predicting aggregation prone hydrophobic regions on the protein surface (Courtois *et al*., 2016; Voynov *et al*., 2009). A development of the SAP software, incorporating charge and hydrophobicity into the developability index, has been reported (Lauer *et al*., 2012). Predicting aggregation risk for antibodies from sequence using bioprocessing data has also been described, with an associated tool available commercially (Obrezanova *et al*., 2015). The freely available CamSol (Sormanni *et al*., 2015, 2017), and Aggrescan 3D (Zambrano *et al*., 2015) servers use sequence and structural information for rational design of mutants with enhanced solubility.

We have recently reported (Hebditch *et al*., 2017a) the protein-sol server (https://protein-sol.manchester.ac.uk/) for sequence-based prediction of protein solubility, calibrated with experimental solubilities in high throughput cell-free expression of *E. coli* proteins. Here, we discuss extension and utility of this freely available web tool, with structure-based calculations. Patch analysis is introduced for electrostatic potential, using Finite Difference Poisson-Boltzmann (FDPB) methods (Warwicker, 1986) that aid visualisation of asymmetric charge distributions. Analysis of non-polar surface uses a patch approach (Cole and Warwicker, 2002), importantly with benchmark analysis of Fab fragments to illustrate the range of values that are associated with surfaces and interfaces. Furthermore, taking into account the common use of pH and ionic strength variation in bioprocessing, a heatmap is produced showing prediction of how protein folded state stability varies with these parameters. Comparison with available data for CH2 and CH3 domains reproduces the qualitative differences observed.

## Methods

### Using the protein-sol patches and heatmap tools

All software at protein-sol is free to use without license or registration and is available online at https://protein-sol.manchester.ac.uk. To use the protein-sol patches or heatmap code, the user simply needs to upload a protein structure in the standard protein data bank format (Berman *et al*., 2000), with results returned in a molecular graphics viewer and as downloadable files, available using a supplied custom URL for 7 days. The protein-sol Webserver is built using open source software. Patches data is displayed using the NGL viewer (Rose *et al*., 2018), and the heatmap visualisations are made in python.

### Protein-sol patches calculation and visualisation

From the supplied PDB structure, only protein is included in the calculation, with the advantage that parameterisation failures for ligands unknown to the dictionary are avoided. Electrostatic calculations follow published protocols from our group (Moutevelis and Warwicker, 2004; Warwicker, 2004), but pKa calculations are not made. Ionisable group charges are fixed at pH 6.3, giving half protonation for histidine sidechains, full deprotonation of aspartate and glutamate sidechains and protein carboxy termini, and full protonation of lysine and arginine sidechains and protein amino termini. Any supplied hydrogen atoms are removed, and polar hydrogen atoms added back, to carry partial charges. Electrostatic potential is calculated with the FDPB method (Moutevelis and Warwicker, 2004) on a 0.6Å spaced grid, with relative dielectric values of 4 for protein and 78.4 for water. Counterions are included at 0.15 M concentration to model ionic strength that matches physiological. For ease of display, both in the server and as a download, the potential is transcribed from the Cartesian calculation grid to the B-factor field of a PDB file containing the original coordinates. To accomplish this with visualisation of potential values at the protein surface, a grid shell surrounding the protein is extracted from the Cartesian grid (Bate and Warwicker, 2004), and potential values assigned to protein atoms according to the closest point on this surface grid shell. Potential values are capped at lower and upper values of −86 and +86 mV, to fit with the PDB B-factor field, these values correspond to an interaction magnitude for a unit charge in the field of about 8.6 kJ/mole. The resulting electrostatic potential surface, and patches, can be manipulated by the user with the NGL viewer (Rose *et al*., 2018). An equivalent colour scheme for potential can also be viewed from the downloadable coordinate file, for example using the red_white_blue spectrum command, with minima and maxima of −86 and +86 in PyMOL (Schrodinger, 2010). In the embedded viewer of the server, various representations other than surface are possible, as are full-screen viewing and picture download.

A different branch of the code evaluates the non-polarity of patches around over the protein surface. For this purpose, a patch is associated with each non-hydrogen atom in the protein. Each patch is the ratio of non-polar to polar solvent accessible surface area for all non-hydrogen atoms within a 13Å radius of the central atom (Cole and Warwicker, 2002). As for electrostatic potential, this property is inserted into the B-factor field of a PDB file, and displayed in the embedded NGL viewer, as well as being downloadable for local viewing. Colour-coding in the embedded viewer is chosen as purple (more polar) to green (more non-polar), so as to distinguish it from the standard red, white, blue scheme for electrostatic potential, and can be visualised using the magenta_white_green spectrum command with minima and maxima of 0.4 and 2.5 in PyMOL (Schrodinger, 2010). It is standard practice to display the most non-polar surface regions, in the context of protein solubility. It is worth noting however that more polar regions also carry information, to our surprise we found that the antigen-combining regions of antibodies are, on average, relatively polar.

### Protein-sol heatmaps for the predicted pH and ionic strength dependence of stability

Whereas the calculation of electrostatic potential on protein-sol is made with standard pKas, it is the differences to standard pKas ΔpKas) that determines the pH-dependent contribution to folded state stability. Not only are pH and ionic strength screens used in formulation studies, but also low pH is used for viral inactivation of biologics expressed from CHO cells (Birch and Racher, 2006). We have developed software in previous applications to compute pKas and the pH-dependent contribution to protein stability, and now provide this code at the protein-sol site. Experimental groups often provide the results of pH and ionic strength screens as heatmaps, and we have therefore chosen this format. Whilst we are unlikely to be describing precisely a feature measured experimentally, folded state stability (for which we provide a prediction of the pH-dependent component) is a key underlying property. Indeed, in the Results section we discuss a qualitative fit between heatmaps generated for CH2 and CH3 domains, and experimental data. Rather than the FDPB model used for electrostatic surface generation, we use the more simple Debye-Hückel (DH) scheme for charge-charge interactions in a medium of uniform relative dielectric (78.4, water) and ionic strength (variable, 0 to 0.3 M). This allows rapid calculation (Warwicker, 1999) of the required Monte Carlo sampling of protonation states (Beroza *et al*., 1991). It is approximated that there are no interactions between ionisable groups in the unfolded state, and the pH-dependent energy is given in Joules per amino acid, a normalisation against protein size. The predicted net charge of the protein (units of e per amino acid) is also given in the heatmap format, within the pH 2 to 8, and ionic strength 0 to 0.3 M ranges. Ligands are, again, excluded from the calculations. In order to give the user context, 2D plots of pH-dependent contribution to stability are drawn for ionic strengths of 0, 0.15, and 0.3 M. The user-supplied protein is displayed against a background of the Fabs dataset analysed in this work.

For calculation with representative CH2 and CH3 domains of an IgG1 antibody, the 1HZH structure was used (Saphire *et al*., 2001). Coordinate files for each domain (CH2 and CH3) were extracted from the overall 1HZH file.

### Datasets of Fab structures for calculation

The Fab dataset was formed by searching the PDB (Berman *et al*., 2000) for structures containing Fabs, and the biological assembly files retrieved. Sequences from the resulting structures were analysed manually to identify only structures with unique heavy and light chains, resulting in 199 Fab structures. From these Fab structures, the four individual domains, the variable and constant domains of the heavy chain (VH and CH), and the variable and constant domains of the light chain (VL and CL), were identified using interdomain sequence motifs (Hebditch *et al*., 2017a). This resulted in 199 Fab structures that constitute the heatmap dataset.

In order to identify antibody:antigen interfaces, coordinates for the antigen binding VH and VL domains were compared with all non-Fab atoms in the relevant PDB file. For any non-Fab coordinate within 10Å of the combined VH and VL domain coordinates, the entire chain of the close non-Fab structure was extracted and combined with the entire Fab, in a new coordinate file. From the original 199 entirely unique Fab structures, 90 of the original biological assemblies contained an antigen within 10Å of the VH and VL domains (Figure 1). These 90 Fabs were then also split into heavy and light chains for chain:chain analysis, and forming the basis for putting the patches part of the server into the context of Fab calculations. We were interested in whether regions of Fab that were not determined to be interfacial (H - L chain or with antigen) were representative more generally of protein surfaces. For this purpose we used a dataset of 54 enzymes (Bate and Warwicker, 2004) known to be monomeric, and thus likely to present mostly non-interfacial amino acids.

**Figure 1:**
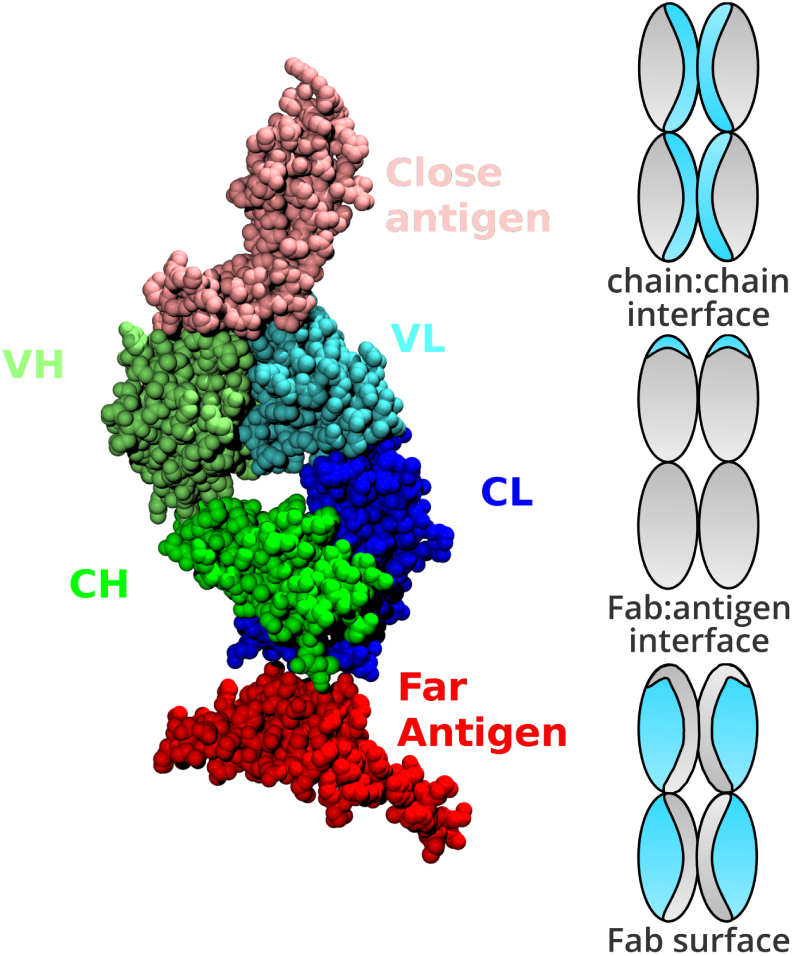
Structural classification of the Fab proteins. On the left, an example of categorisation of chains in a Fab PDB file. Although each Fab may contain multiple non Fab chains, we elected to only consider chains within 10Å of the Fab VH and VL domains as the antigen. As a result, the Fab:antigen interface is that between the VH and VL of the Fab, and any non-Fab chain within 10Å. On the right, a schematic representation of classification for atoms in a Fab fragment. Regions highlighted in blue denote interface (top and middle) or surface (non-interface, bottom). Assignment to the different categories was made from calculations of SASA, as described in the text.

## Results

### Categorisation of surface, buried and interface atoms

Solvent accessible surface area (SASA) was calculated for each atom in each construct: the extracted Fab alone, the extracted antigen alone, each Fab chain alone, and the Fab:antigen complex. With solvent accessible surface areas for each atom in each construct ascertained, we then assigned structural categorisations (Figure 1) based on solvent accessible surface area. An atom was defined as buried for SASA < 5Å, and surface accessible otherwise. A lower threshold (0.1 Å) was used to assess change in SASA for an atom, upon interface formation, and assign to the relevant interface (Fab:antigen or Fab chain:Fab chain). Once each atom is tagged with one or two of the above three tags, it was assigned a single structural categorisation, prioritising the interface over surface categorisation. Thus, atoms with a surface categorisation are at the Fab surface and outside of both interface types (Fab:antigen and H chain: L chain). As a result it is possible, for a dataset of Fab fragments, to compare the two interface environments and the remaining surface regions. The ratio of non-polar to polar SASA (NPP) in an interface are assigned from the constituent parts of the complex that contains that interface. For example, for Fab:antigen, Fab fragment NPP ratio values are taken from the Fab calculation. It is then possible to compare the distributions of NPP ratio for interfacial (including different types of interface) and surface atoms. A similar comparison can be made for distributions reduced to just the set of maximal NPP ratio values, where a maximum is taken from each environment (interface, surface) in each Fab system. For further comparison, the surfaces of a monomeric enzymes set are also included, presumably representative of few interacting surfaces.

### One step visualisation of charged and hydrophobic surface patches

The protein-sol patches software takes a protein data bank (PDB) structure and calculates patches of charge (at pH 6.3) and hydrophobicity across the protein surface (Figure 2). This allows the user to quickly identify interesting regions on the protein surface which may influence the behaviour and stability of the protein structure. Electrostatic surface potential based on FDPB calculation is plotted alongside the potential colour-code. Note that a relatively large change in pKa for acidic and basic groups in a surface salt-bridge may be about 1 pH unit, equivalent to 57 mV i.e. the range of potential given here is that expected for electrostatic interactions at the protein surface. Potential equivalent to thermal energy at 300 K, kT/e, is 25 mV. A scale is also given for colour-coding by patch NPP ratio, from 0.6 (more polar) to 2.5 (more non-polar). Importantly, an additional bar graphic displays the maximum of NPP ratio, in the context of maxima found for interface and non-interface regions of Fab fragments. This information allows the user to find not just the most non-polar region, but also to assess its significance against known interfaces, significantly enhancing the practicality of the tool.

**Figure 2:**
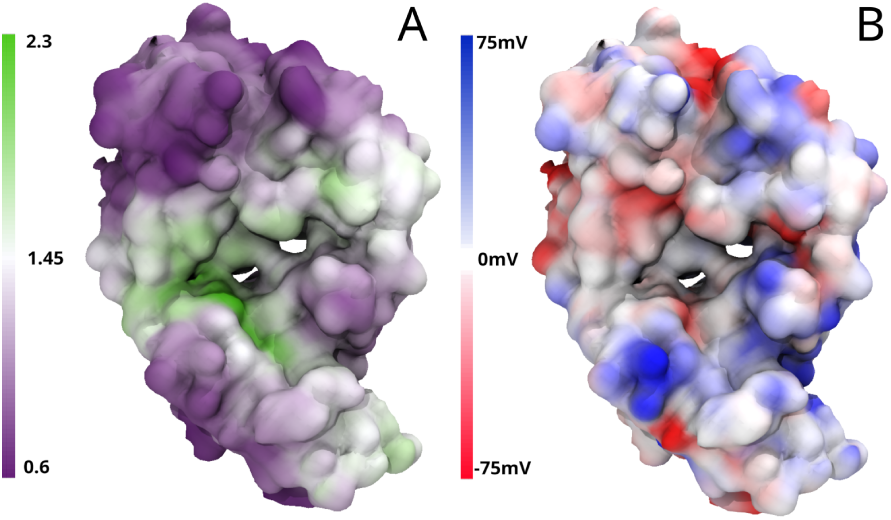
Example protein-sol visualisation of surface patches on a Fab. In A) the Fab is colour-coded from low NPP ratio (purple) to high NPP ratio (green), and in B) the Fab is colour-coded from negative charge (red) to positive charge (blue). Both are visualised using the embedded NGL viewer on the protein-sol web application after calculation.

Using our dataset of different Fab structural categorisations, we demonstrate the potential of the protein-sol patches software to quickly identify hotspots of relative hydrophobicity (higher NPP ratio). Through interrogation of the dataset for extremes, Figure S1 shows particularly hydrophobic patches for the Fab chain:chain interface (Figure S1A), a patch on an interface between Fab fragment and antigen (Figure S1B), and a patch on the surface of a Fab fragment (Figure S1C). Note that these are extreme values and, as we show in a subsequent section, antibody-antigen interfaces are, on average, relatively polar.

### Heatmaps show the predicted pH and ionic strength dependence of stability

Alongside surface visualisation, protein-sol also provides heatmaps for the pH and ionic strength dependence of folded state protein stability, using the Debye-Hückel (DH) method for interactions between ionisable groups, and pKa calculations. Two separate heatmaps (Figure 3) display predicted charge (units of e per amino acid), and predicted pH-dependent contribution to stability (J per amino acid). Normalisation relative to sequence length allows direct comparison of proteins. Each heatmap consists of 91 combinations of pH and ionic strength. In order to compare qualitatively with experiment, CH2 and CH3 domains from the IgG1 PDB structure 1HZH are used (Saphire *et al*., 2001), since pH and ionic strength variations in stability have been reported for IgG1 CH2 and CH3 domains (Yageta *et al*., 2015). Measured phase diagram boundaries for these domains, derived at acidic pH (Yageta *et al*., 2015) has been marked (Figure 3), showing a good match between these calculations and experiment. For example, looking across the pH range at 0.15 M ionic strength, there is a greater variation in pH-dependence for CH2 than for CH3 domains. The pH-dependence of stability is directly related to charge (Antosiewicz *et al*., 1994), but perhaps the most convenient feature to extract from heatmaps of charge is the predicted sign of net charge. It should be noted that these calculations lack post-translational modifications, such as glycosylation. Here, the IgG1 CH2 domain is predicted to be rather more positively-charged than the CH3 domain at equivalent pH and ionic strength.

**Figure 3:**
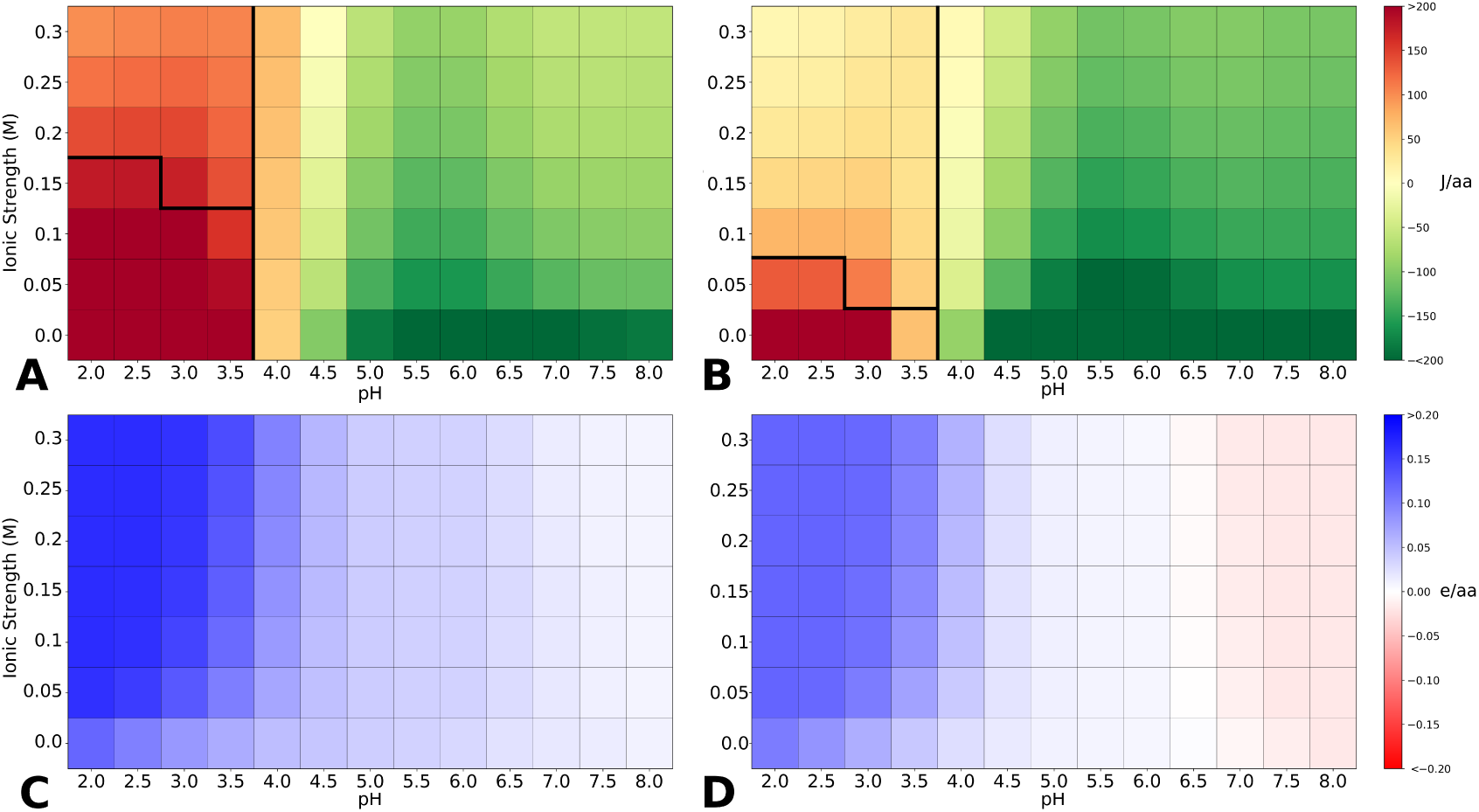
Protein-sol stability and charge heatmaps. Calculated folded state stability for the CH2 (A) and CH3 (B) domains, and charge heatmap calculated for the CH2 (C) and CH3 (D) domains of a monoclonal antibody (1HZH).

Differences in the physicochemical screens represented by the heatmaps can also be visualised in terms of line graph streams that show the variation with pH of charge or energy at a fixed ionic strength (Figure S2). Our dataset of Fab fragments is used as a background with which to compare the pH dependence calculation for the user structure. We chose this background set due to its importance in the biopharmaceutical and biotechnology areas. Wider-scale analysis is possible, but analysis of proteins that are native to differing environments can be a complex topic (Chan and Warwicker, 2009). Comparison of Figure S2 panels A and B show clearly that the predicted pH-dependence of stability is greater for the CH2 domain for the CH3 domain. Similarly, Figure S2 panels C and D show the more positive predicted charge of the CH2 domain relative to the CH3 domain, at equivalent pH values.

### A common distribution of patch polarity for protein surfaces

To provide the user with context for patch polarity, alongside surface display, it was necessary to undertake a bioinformatics analysis. Protein surfaces were studied for 3 datasets, the Fab fragments, their corresponding (protein) antigens, and a set of enzymes that are monomeric in their biological states. Surfaces were assigned for Fab fragments and antigens, excluding interfaces, as described in the Methods section, and the entire surfaces of the monomeric enzymes were included. The distribution of NPP ratios are similar for all 3 datasets (Figure 4A), giving confidence that this form of distribution is broadly representative of protein surfaces. Since a higher NPP ratio reflects a more hydrophobic patch, and a lower NPP ratio relates to a more polar patch, the similarity in distributions suggests that the polar to non-polar spectrum of a protein surface is a general property. All three protein sets have a peak in the distribution at an NPP ratio around 1.0, i.e. with equal contribution from polar and non-polar surface areas.

**Figure 4:**
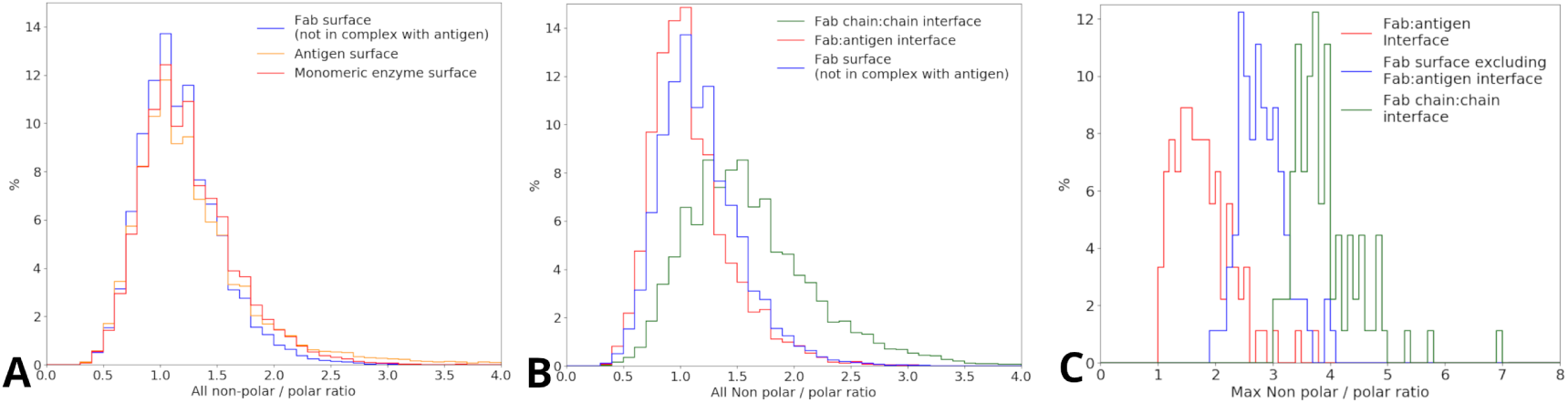
Distribution of hydrophobic patches across the different Fab structural classifications. A) Distributions of all NPP ratio values for the Fab, antigen and monomeric enzyme surfaces combined for each protein. B) Distribution of all NPP values for all three Fab structural categorisations: Fab:antigen interface, Fab surface (excluding Fab:antigen interface) and Fab chain:chain interface combined for each protein and compared. C) Distribution of max NPP ratio values for Fab:antigen interface, Fab chain:chain interface, and Fab surface.

### The heavy to light chain interface is more non-polar than protein surface, but the Fab-antigen interface is relatively polar

Having ascertained that the Fab protein surface is representative of protein surfaces in general, a comparison is made with heavy to light chain and Fab-antigen interfaces (Figure 4B). As expected for a protein-protein interface, the NPP ratio distribution is substantially shifted towards non-polar for the chain-chain interface within a Fab fragment, By comparison, the Fab:antigen interface is slightly more polar than the Fab surface, suggesting that the two interface types have different physicochemical properties. The polarity surface displayed on the server relates to the NPP ratio distributions given in Figures 4A and 4B. In order to give the user a more succinct indication of where a protein fits in terms of non-polar surface, we decided to extend the analysis to record the patch with the highest NPP ratio, in a particular protein. It is this property that is shown in the bar chart displayed following the surfaces on the server. Figure 4C shows distributions of the highest NPP ratio values, extracted (one for each protein) from the full distributions (Figure 4B). Interfaces within a Fab fragment are again, on average, shifted towards more non-polar. The relative polarity of Fab-antigen interfaces is now even more apparent, clearly shifted towards more polar than Fab surface. For each Fab, the most hydrophobic patch is generally in the the Fab chain:chain interface, and the least hydrophobic in the Fab:antigen interface. Overall, this approach allows a user to find the single largest non-polar patch within a structure, and is typically related to features used in the assessment of the developability for biotherapeutics.

## Discussion

The development and use of therapeutic proteins can be limited by instabilities which complicate manufacture, storage and delivery. It is important to improve understanding and to provide predictions for the factors that cause reversible and irreversible association. To help improve the developability of biopharmaceuticals, in past work, we introduced the protein-sol sequence software for predicting protein solubility based on primary structure (Hebditch *et al*., 2017a). In this work, we introduce two new tools, available for free and with no licensing requirements, protein-sol patches and heatmaps. Whilst targeted at the biopharmaceutical research community, they could also be of wider interest for biotechnology. Both hydrophobic and charged surface patches have been implicated in aberrant solution behavior. The protein-sol patches code is used to calculate the predicted surface patches from a protein structure. Incorporation of the NGL viewer allows fast and simple visualisation of the surface electrostatic potential and polarity (hydrophobicity). Calculation results are also available for download, as PDB format files with surface patches colour-coded using the B-factor field. Results are therefore readily available for further processing or visualisation by the user.

To put the server output into context, we investigated the hydrophobic properties of surface and interfacial regions of Fab fragments, as well as a dataset of soluble monomeric enzymes. There is little difference in NPP ratio distributions for surface regions of Fab fragments, their corresponding protein antigens, and monomeric enzymes (Figure 4A). One interpretation of this result is that a standard protein surface profile of polarity is associated with a balance between structural stability and solubility. As expected, the heavy - light chain interface is relatively non-polar, both in overall distribution (Figure 4B) and as peak values for each Fab (Figure 4C). Surprisingly though, Fab-antigen interfaces are relatively polar, an observation that is exaggerated when viewed as peak NPP ratios (Figure 4C), as compared with the whole distributions (Figure 4B). Antibody-antigen interfaces have been reported to differ from other protein-protein interfaces, tending to be smaller in size, incorporating fewer helices and more loops, with less hydrophobic packing (Dalkas *et al*., 2014). We now find that antibody-antigen interfaces are, if anything, even more polar than surface (non-interfacial) regions. It is possible that the constraints of altered interfacial size and different secondary structure composition lead to a reduction in non-polarity. An alternative possibility is that as a non-obligate interface, there remains a requirement for solubility in the absence of interface formation. Further, following reports that proteins at high naturally occurring concentrations tend to more soluble (Tartaglia *et al*., 2007; Hebditch *et al*., 2017b), the constraint for relatively polar surface would be enhanced for antibodies. The observation of relatively polar antibody-antigen interfaces is not necessarily inconsistent with the finding that *π-π* interactions are common (Dalkas *et al*., 2014). Differences were computed between amino acid percentage compositions overall in Fab fragments and specifically in CDRs, for the dataset used in this study. The largest such difference is for tyrosine, at 5.9%, whilst the other aromatic sidechain amino acid differences are much lower, at 1.5% (phenylalanine) and 0.4% (tryptophan). Next largest after tyrosine is valine (−4.4%). Thus some compensatory effect may exist, with elevation of tyrosine (and the potential for *π-π* interactions) countered by lowering of hydrophobic valine sidechains.

The NPP ratio part of the protein-sol server has been developed to allow users to view nonpolar patches in the context of developability. Incorporation of modal values from the NPP peak values distributions in Figure 4C, alongside the NPP peak value for the user’s protein in a simple graphic, will aid such assessment. Whilst non-polar patches may be the focus for developability, we do not discard more polar regions from the display. A demonstration of the relative polarity of antibodies at the antigen-combining site is shown in Figure 5. Whereas for Figure S1B we chose the most non-polar complementarity-determining region (CDR) (with a green patch), more typically the relative polarity is apparent (Figure 5). Indeed, since this graphical tool makes the separation of relatively polar and non-polar regions readily apparent, it could accelerate discoveries such as that made here for antibody - antigen interfaces. The composition of F,W,Y amino acids in the CDRs of the Fab shown in Figure 5 (3mly) is 14.8%, within a range of 10% to 29% for CDRs in the set of Fabs used. By contrast, the F,W,Y composition for the entire Fabs is 8.7% for 3mly, and range from 7% to 12% for the set. Our analysis of surface non-polarity with a 13Åradius patch does not, in general, highlight CDR surfaces. Figure S3 shows the effect of reducing patch radius for 3mly, non-polar patches do now appear within the CDRs, but these remain smaller non-polar features than seen elsewhere on the Fab surface. We infer that the use in CDRs of amino acids with aromatic sidechains is more subtle than can be judged simply by scale of exposed non-polar surface area.

**Figure 5:**
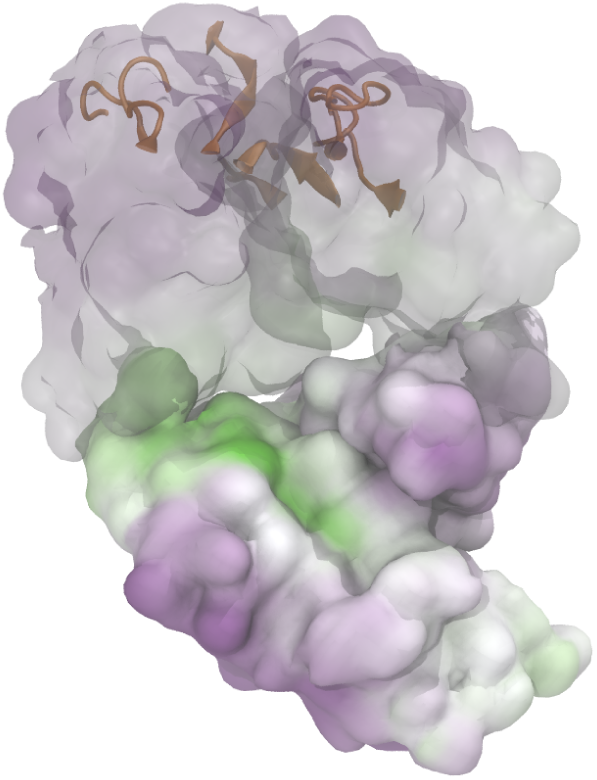
Example visualisation of the relatively high polarity at the Fab antigen interface in comparison to the rest of the Fab. Regions of high hydrophobicity are coloured green, low hydrophobicity coloured purple, and CDR regions highlighted in orange.

The current work provides an additional tool for groups looking to identify regions of a protein for engineering improved solution properties. Both hydrophobic patches (Nichols *et al*., 2015), and charged patches (Yadav *et al*., 2012; Chow *et al*., 2016; Perchiacca *et al*., 2012) have been mutated to alter solution behavior. The ability to rapidly visualise surface patches will further inform and accelerate such work.

With regard to heatmap depiction of predicted stability for pH and ionic strength variation, these are two important factors when studying proteins. Although the trend of changes will be uniform, acidification tends towards a more positive protein and increased ionic strength reduces electrostatic interactions, the net outcome is a delicate balance of the constituent parts. For example, we demonstrate (Figure 3) that qualitative experimentally-determined differences between IgG1 CH2 and CH3 domains (Yageta *et al*., 2015), are reproduced by our calculations. Furthermore, the user can view the size of predicted pH-dependence as a comparison with the dataset of Fab fragments, with plots normalized for sequence length. Predicted protein charge is also presented, in an analogous manner. Fab fragments were used as the control dataset since they are a widely used biopharmaceutical platform. We suggest that the protein-sol heatmap may be a useful tool for accelerating formulation screens by identifying potentially favourable regions prior to formulation development, when a structure or structural model is available. Since viral clearance procedures often involve a low pH step, the heatmap analysis will aid determination of the degree to which protein stability is diminished as salt-bridges and other favourable interactions are lost at acidic pH.

In this work we have discussed how the polar, non-polar, pH and ionic strength dependent properties influence protein stability in solution, and how instability can limit development. The protein-sol software suite, incorporating patches and heatmap software has been designed to benefit researchers interested in understanding the surface properties, and stability, of proteins in solution. While developing the server, we have demonstrated how the patches software can identify interesting physicochemical properties of Fab chain:chain and Fab:antigen interfaces, and also how predictions for protein stability compare favourably to measured data. Protein-sol is freely available. Our initial work suggests that it could contribute to the acceleration of protein engineering and formulation optimisation, and to the improvement of developability for new biotherapeutic leads. It also provides insight into the fundamental properties of proteins in solution.

## Data Availability

The datasets analysed during this study are available from the corresponding author on reasonable request. The reported web tool is freely available online.

## Acknowledgements

Members of the Curtis and Warwicker groups are thanked for discussion and providing feedback. The authors would like to acknowledge the assistance given by IT Services at The University of Manchester.

## Author contributions statement

Both authors conceived and conducted the experiment(s), analysed the results and reviewed the manuscript.

## Additional information

### Competiting interests

The authors declare that they have no competing interests.

